# Non-Negative Matrix Factorization for Analyzing State Dependent Neuronal Network Dynamics in Calcium Recordings

**DOI:** 10.1101/2023.10.11.561797

**Authors:** Daniel Carbonero, Jad Noueihed, Mark A. Kramer, John A. White

## Abstract

Calcium imaging allows recording from hundreds of neurons *in vivo* with the ability to resolve single cell activity. Evaluating and analyzing neuronal responses, while also considering all dimensions of the data set to make specific conclusions, is extremely difficult. Often, descriptive statistics are used to analyze these forms of data. These analyses, however, remove variance by averaging the responses of single neurons across recording sessions, or across combinations of neurons, to create single quantitative metrics, losing the temporal dynamics of neuronal activity, and their responses relative to each other. Dimensionally Reduction (DR) methods serve as a good foundation for these analyses because they reduce the dimensions of the data into components, while still maintaining the variance. Non-negative Matrix Factorization (NMF) is an especially promising DR analysis method for analyzing activity recorded in calcium imaging because of its mathematical constraints, which include positivity and linearity. We adapt NMF for our analyses and compare its performance to alternative dimensionality reduction methods on both artificial and *in vivo* data. We find that NMF is well-suited for analyzing calcium imaging recordings, accurately capturing the underlying dynamics of the data, and outperforming alternative methods in common use.

## Introduction

Recent advances in neural recording methods have made it possible to collect large, complex data sets that can be used to study context-dependent neural dynamics in great detail ^1,2^. For example, calcium imaging allows simultaneous recording from hundreds of neurons in a single brain region *in vivo*, while maintaining single cell-resolution ^3,4^. Other tools allow cell “tagging” based on factors, including activity during a specified time window and molecular genotype ^5–8^. The resultant data set consists of several hundred time series thousands of points long, multiplicatively increasing per experimental condition added, in addition to any tagging data. The process of characterizing such single-cells responses, relating them to those of other cells across time, and accounting for the tagging information, all while considering changes in animal behavior, is a difficult but essential task.

Basic descriptive statistics (mean, median, variance) are a common approach to analyze large neural datasets. However, these approaches are limited because they average over the temporal dynamics of neuronal activity and can obscure interactions among neurons at fine time scales ^9,10^. Dimensionality reduction (DR) is a more sophisticated approach that can help to reveal low-dimensional dynamics by transforming high-dimensional data into interpretable, low-dimensional components ^11^. Compressing the data in this way can preserve temporal patterns and correlations and focus subsequent analysis on the dynamical patterns of interest.

DR approaches have many applications in the analysis of neural data, including analysis of electrophysiological recordings ^9,12,13^, automation of trace extraction of neuronal activity in fluorescent recordings ^1,14–18^, and as a pre-processing step to more complex behavioral decoding ^19^. To analyze network dynamics in calcium recordings, non-linear DR methods have emerged to map dynamics to low dimensional manifolds ^20–23^. While excellent for very low dimensional visualization ^24,25^, the dynamics are difficult to interpret beyond the manifold being modeled, with manifolds being difficult to interpret beyond three dimensions. Further, dynamical non-linear methods have been applied to recover neural trajectories as a function of latent variables^26^. Linear DR methods have likewise been employed to either statically cluster cells ^18,27–29^, or to abstractly analyze activity in low-dimensional subspaces ^30–33^.

While each methods have their own strengths as a function of different low dimensional representations, we aim to decompose our data into a lower dimensional series of neuronal sub-networks and their activities, to detect how neurons form flexible, correlated cell assemblies under different experimental contexts. We argue more generalized linear DR methods will provide a more interpretable representation of the data, at the cost of a few more dimensions than non-linear methods, dynamical or non-dynamical, as a function of having fewer tunable parameters and simpler mathematical decompositions. We further argue Non-negative Matrix Factorization (NMF) is an especially promising linear decomposition method for analysis of activity recorded in calcium imaging in this context. NMF operates under the constraint that every element in the data, and the decomposition, be non-negative ^34^, as is naturally true for neuronal calcium traces ^18^. Further, the positivity constraint, coupled with the linearity of the reduced dimensions, ensures that each of the dimensions sum to form a low dimensional representation of the data, simplifying the interpretation of the data ^35^. In neuroscience, specifically in the context of calcium imaging, NMF has emerged as the state of the art method to estimate single neuron activity via fluorescent recordings ^14–16^. However for analyzing the dynamics of recorded neurons, it has been applied sparingly, either for quantifying single cell pairwise relationships ^36^ or statically clustering cells under a series of strict assumptions ^28^. Here, we develop an application of NMF to analyze holistic network activity, comparing it to the most widely employed methods that provide a similar decomposition, capturing how precise sub-networks of neuronal dynamics evolve in different experimental contexts. To assess performance of the NMF approach, we generate artificial calcium imaging data from simulated ground-truth networks and assess how well our approach captures the underlying network structures responsible for the emergent dynamics of the network. We then apply the same NMF framework to characterize the dynamics of a representative *in vivo* calcium imaging data set. We conclude that, compared to alternative methods, NMF best captures the ground truth networks that underlie the observed calcium dynamics and provides a useful low-dimensional description of complex physiological data.

## Dimensionality Reduction (DR) Methods

### Non-Negative Matrix Factorization (NMF)

NMF is a linear, matrix-decomposition method requiring all elements be non-negative ^34^. Each reduced dimension, or component, identified in NMF can be interpreted as a specific combination of input features representing distinct sources of variance in the data ^35^. The components then sum to represent the original data, rather than canceling variables (using negativity) to achieve a geometrically strict representation ^37^. Mathematically, NMF decomposes an original matrix X into two constituent, lower rank, matrices W and H, given by:

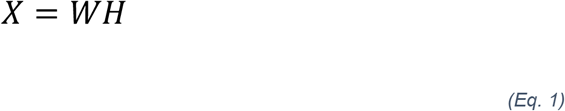

with the aim of iteratively updating W and H to minimize an objective function so that the product of the two deconstructed matrices optimally reconstructs the original ^34^. Here, X is a column-wise representation of the data, where each column is the time-series fluorescent activity of a neuron (Fig. 1b) extracted from the recording (Fig. 1a), that is decomposed into two lower rank matrices, W and H. For NMF, W can be interpreted as a feature matrix, each vector representing a distinct pattern in the input data, and H interpreted as a coefficient matrix, representing the contribution of each feature vector to each point in the original matrix. When using DR for analyses, especially in the context of neural recordings, the underlying assumption is that the original matrix can be decomposed into a lower rank series of matrices. The rank, k, should be significantly smaller than n (k << n), the number of neurons recorded, while still recapturing a majority of the variance present in the data ^38^. Model performance can be assessed by the variance explained as represented by the coefficient of determination (R^2^) ^39^, given by:

**Fig. 1.**
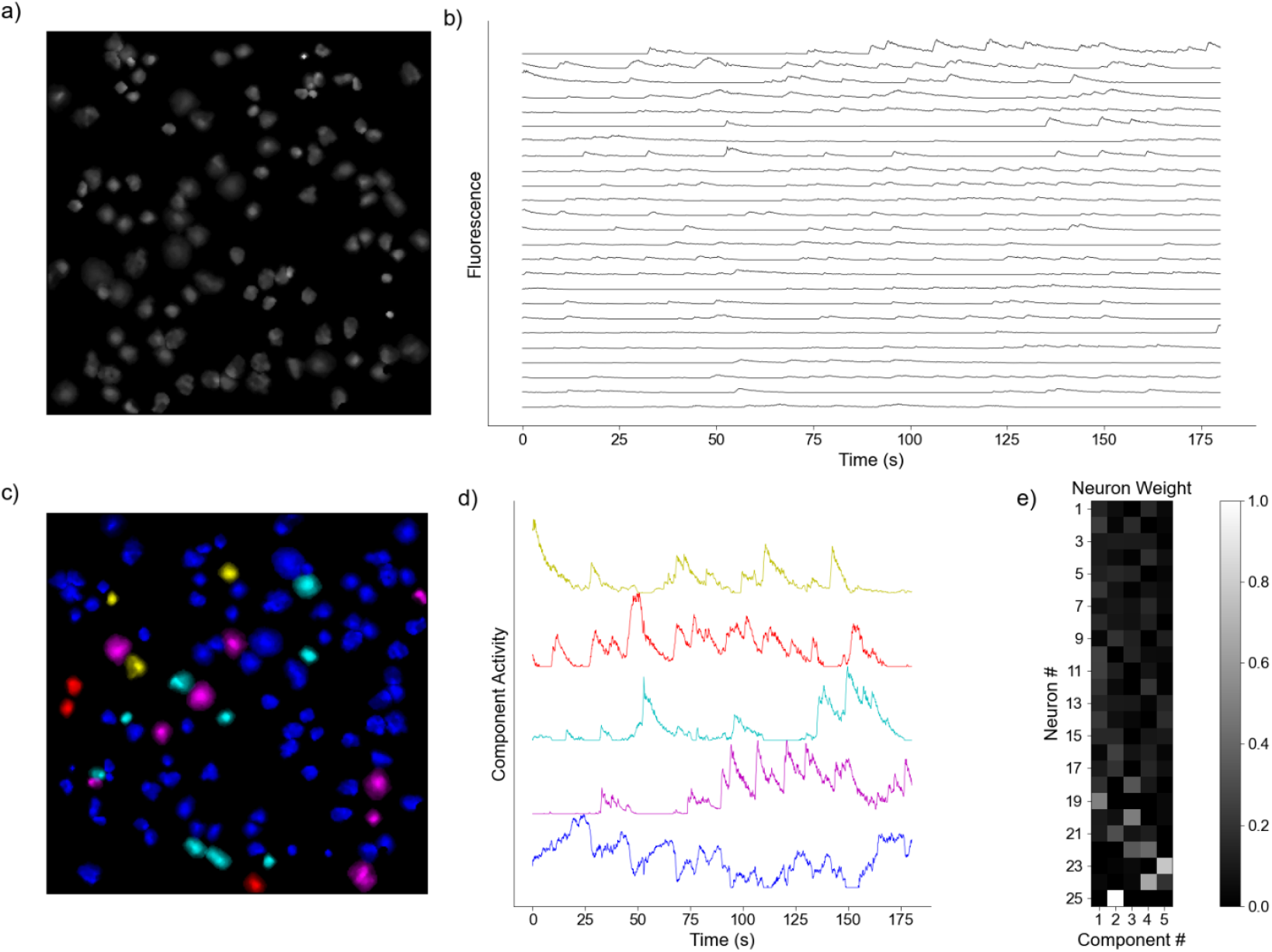
Representative Model Pipeline. **a)** Recording Field of View (FOV) with Regions of Interest (ROIs). **b)** Extracted fluorescent activity traces from ROIs. **c)** Same recording FOV shown in a) with ROIs colored according to largest component contribution. **d)** Decomposed activities of the components shown in c). **e)** Sample truncated heatmap showing neuronal contribution or weight for each component

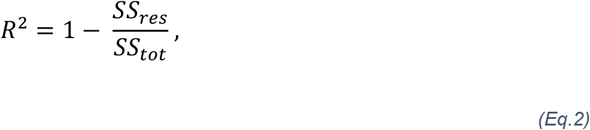

where,

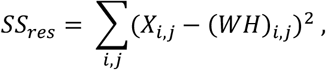

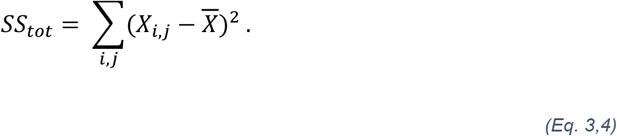

Thus, the model captures the original data perfectly in the limit as R^2^ approaches one. R^2^ increases as components are added, because the rank of the two constituent matrices are increasing. Akaike Information Criterion (AIC) has been adapted for NMF to inform the rank to which the dimensionality should be reduced ^35,40^, with the optimal model minimizing it ^41^; for our implementation of NMF:

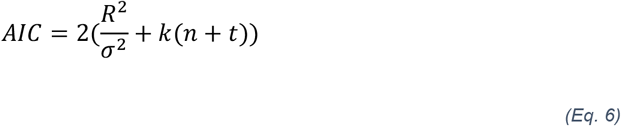

Where R^2^ is the reconstruction error defined in Eq. 2, σ^2^ is the estimated variance of the data set, k is the number of components the model is decomposing into, n is the number of neurons in the recording, and t is the number of time points in each of the time series being decomposed. For our analysis, we interpret each row of H as a sub-network, with each entry of that row as a neuron’s contribution of to that sub-network (Fig 1e). Each column of W is then interpreted as the time-series for that sub-network (Fig 1d). Further, the neurons in the original recording FOV can be artificially reconstructed to provide visualization of sub-network contribution (Fig 1c). We provide the pseudocode from an implementation of the NMF analysis pipeline:

Input: X [time points x neurons]

Output: NMF_model

1. normalize data between (0,1)*
2. initialize list of AICs
3. for i in range(1, num_neurons)
4. W, H = init_WH(n_components = i)
5. NMF_model = optimize(X, W, H)
6. AIC_list.append(NMF_model.aic)
7. if AIC[i-1] < AIC[i]:
8. n_components = i-1
9. break
10. W,H = init_WH(n_components = n_components)
11. NMF_model = optimize(X,W,H)

* we normalize using the global maximum and minimum of the trace set to preserve the relative difference between them

The above pipeline describes the sequence of using AIC to find an optimal number of components, and then fit a model to that number of components. Our implementation of NMF is a modification of the implementation found in Python’s Scikit-Learn package ^42^. We manually initialize W and H, as opposed to the automated method found in the package, due to the wide variety of initialization methods possible for NMF ^43,44^, and to maintain consistent, precise, and easily reproduceable control over the initial conditions of our models. For the analyses found in this paper, we apply non-negative dual singular value decomposition (nndsvd), a consistent and efficient initialization method ^45^. Further, we implement variance explained and AIC for automatic model selection, as they are not part of the base implementation in the package.

### Principal Component Analysis (PCA)

We implement PCA because of its widespread use in neural data analysis ^12,18,30–32,46–48^. PCA, efficiently calculated by a Singular Value Decomposition (SVD)^37^:

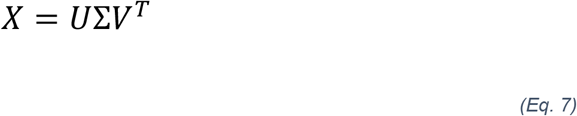

where X is a column-wise representation of our data (i.e., each column is the time-series fluorescent activity of a neuron, assumed to have zero mean) decomposed into U, Σ, and V^T^. V^T^ is the set of basis or singular vectors, also referred to as the principal components (PCs) of the original matrix, where each row of this matrix is a component of the data. Further, each of the PCs are necessarily orthogonal to each other by construction. U is the projection of the original matrix X onto the principal components in V^T^, and Σ is a matrix containing the singular values of X describing the magnitude of the transformation, ordered by decreasing singular values ^37^. The variance explained by a PC is the proportion of its singular value compared to the sum of all singular values. For our analysis, we interpret each PC as a sub-network or prevalent pattern of activity, with each entry of the PC being a neuron’s weight or contribution to that PC. We then interpret the projection of the data onto each PC as the activity or time series of that sub-network, attributing the dominance of that sub-network to the variance explained. Our implementation of PCA is a modification of the implementation found in Python’s Scikit-Learn package ^42^. We provide the pseudocode from an implementation of the PCA pipeline below:

Input: X [time points x neurons] Output: PCA_model

1. normalize data between (0,1)*
2. subtract mean of each time series
3. PCA_model = PCA(X) #data must be in column-format at this step.

* we normalize using the global maximum and minimum of the trace set to preserve the relative difference between them

This is a modification of the implementation of PCA found in sklearn ^42^. PCA calls for every variable, or trace in our case, to be centered about zero^37^. However, we remove the uniform normalization from sklearn’s implementation to allow for finer tuning of data normalization before decomposition with the model.

### Independent Component Analysis (ICA)

We implement ICA because of its ability to extract independent sources from complex signals, which could help separate overlapping patterns of neural activity in a single calcium recording. While ICA has been used for automated trace extraction in calcium recordings ^49^, this approach has seen limited application in the further analyses of neural signals estimated calcium recordings. ICA is a statistical technique that aims to decompose a multivariate signal into statistically independent components ^50^, rather than a set of orthogonal components, as found in PCA. ICA operates under the assumption that k independent sources produce the data, and that the data are a linear mixture of these underlying sources. ICA assumes the data to be a linear mixture of sources:

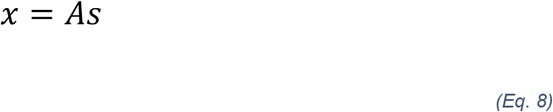

where x is the original data, s are the underlying independent sources, and A is a mixing matrix that mixes the components of the sources ^50^.

In practice, the independent sources found in s, are decomposed using an unmixing matrix, W, such that the linear transformation of the data by W, estimates the underlying independent sources ^50^:

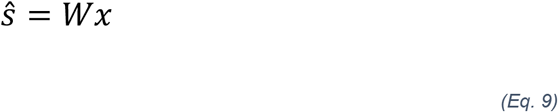

Given that W transforms x to estimate the independent sources, we interpret the components of W as the contribution of each neuron to the sub-network activities, which are the independent sources in s. Here, we interpret the s independent sources as the underlying patterns of activity of each sub-network that drive the observed dynamics. Like PCA, ICA assumes that inputs are zero mean. While PCA is designed to maximize the variance of the data along the principal components ^37^, ICA attempts to separate the data into k independent sources. We use the fastICA implementation found in sklearn ^42^, and provide the pseudocode from an implementation of the ICA analysis pipeline below:

Input: X [time points x neurons]

Output: ICA_model (contains A, s, amongst other information)

1. normalize data between (0,1)*
2. subtract mean of each time series
3. ICA_model = ICA(X)

* we normalize using the global maximum and minimum of the trace set to preserve the relative difference between them

While ICA calls for unit variance for each of the features, which in this case would be the calcium time series, we maintain the relative difference in variance between each neuronal time series to maintain the relative difference in activity recorded as a function of the fluorescence.

### Uniform Manifold-Approximation Projection (UMAP)

We implement UMAP because of its increasing use to analyze life science data ^51,52^ and its ability to capture non-linear relationships. UMAP has been shown to capture complex patterns to visualize and cluster data in very low dimensions ^53^, though the technique remains untested on calcium recordings. Briefly, UMAP aims to optimize between preserving local and global structures, where structure is local between neighboring points and global between data points extending beyond the neighborhood. In neuronal calcium imaging, “neighborhood” refers to local relationships among neurons with similar activity patterns at a time point, while global relationships extend beyond those neighbors at the same point. UMAP subsequently directly maps this structure to a lower dimension ^53^, resulting in the output being a single matrix (the equivalent of H, V^T^, and S), and not a product of matrices. Therefore, UMAP does not provide temporal information about the activity of the lower dimensional patterns or sub-networks (Table 1). Further, there is no well-established way to quantify variance explained in UMAP as in the other linear methods. We use the python package UMAP-learn to implement UMAP, and provide the pseudocode from an implementation of the UMAP analysis pipeline below:

**Table 1.**
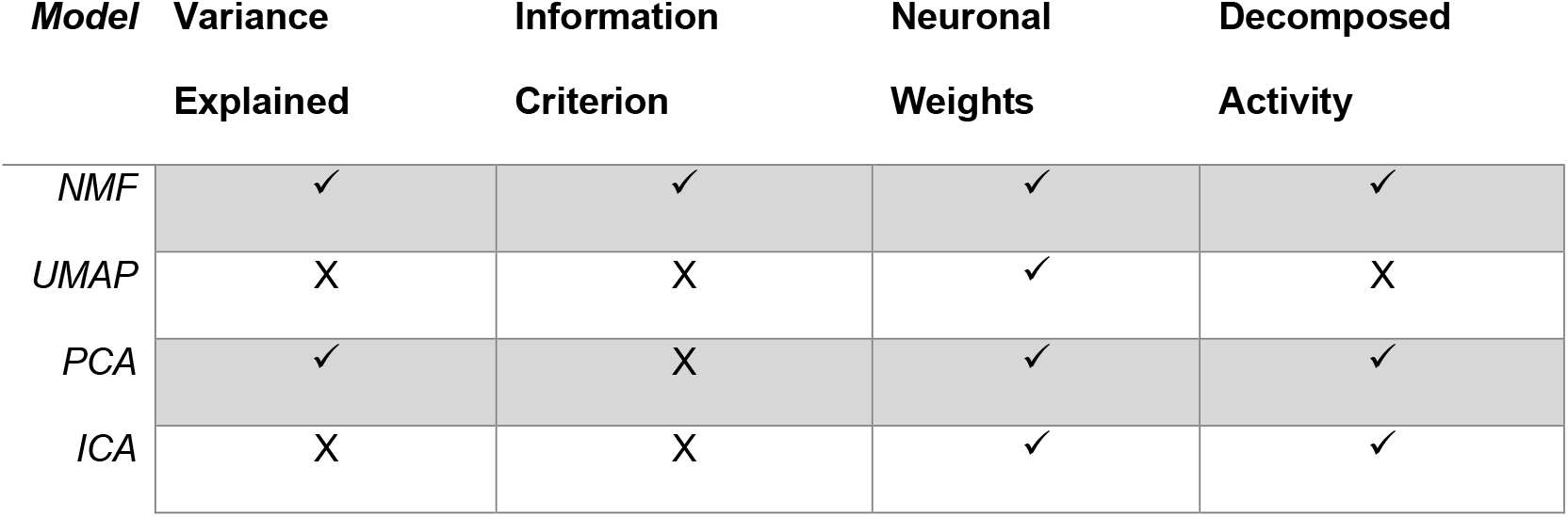
Model Attributes

| **Model** | **Variance Explained** | **Information Criterion** | **Neuronal Weights** | **Decomposed Activity** |
| --- | --- | --- | --- | --- |
| *NMF* | ✓ | ✓ | ✓ | ✓ |
| *UMAP* | X | X | ✓ | X |
| *PCA* | ✓ | X | ✓ | ✓ |
| *ICA* | X | X | ✓ | ✓ |

Input: X [neurons x time points] (NOTE TRANSPOSITION FOR THIS CASE, COMPARATIVELY) Output: UMAP_model (contains low_d_projection, amongst other information)

1. normalize data between (0,1)*
2. UMAP_model = UMAP(X)

* we normalize using the global maximum and minimum of the trace set to preserve the relative difference between them

In addition to the number of components, k, being a manually set hyperparameter, the number of points in the neighborhood is also manually set, in addition to the other 37 tunable parameters in the package ^53^. Because our main objective is to identify how neurons group to form sub-networks and shift as a function of experimental contexts, we apply UMAP to decompose the data X into a *k x n* matrix. k is the number of components or subnetworks we determine to decompose into, while n is the number of neurons, aiming to give a similar representation to H, V^T^, and S in our other methods.

## Network Simulation Methods

To assess performance of the DR methods, we implement two simulation paradigms. In each paradigm, we simulate the spiking activity in a network of interconnected neurons and estimate calcium imaging traces for each neuron. We then apply each DR approach to the simulated calcium fluorescence data. We describe each simulation paradigm below.

### Perfectly Intraconnected, Independent Nodes

We first consider a network of 100 neurons organized into 5 independent nodes, each consisting of 20 neurons. Neurons within the same node are perfectly connected, such that a spike by any neuron in one node produces a spike in all other neurons in that node. Neurons in different nodes are disconnected, such that activity in two neurons of different nodes are independent (Fig. 2a). The activity of a single neuron is governed by a basic Poisson spike generator ^54^, given by:

**Fig. 2.**
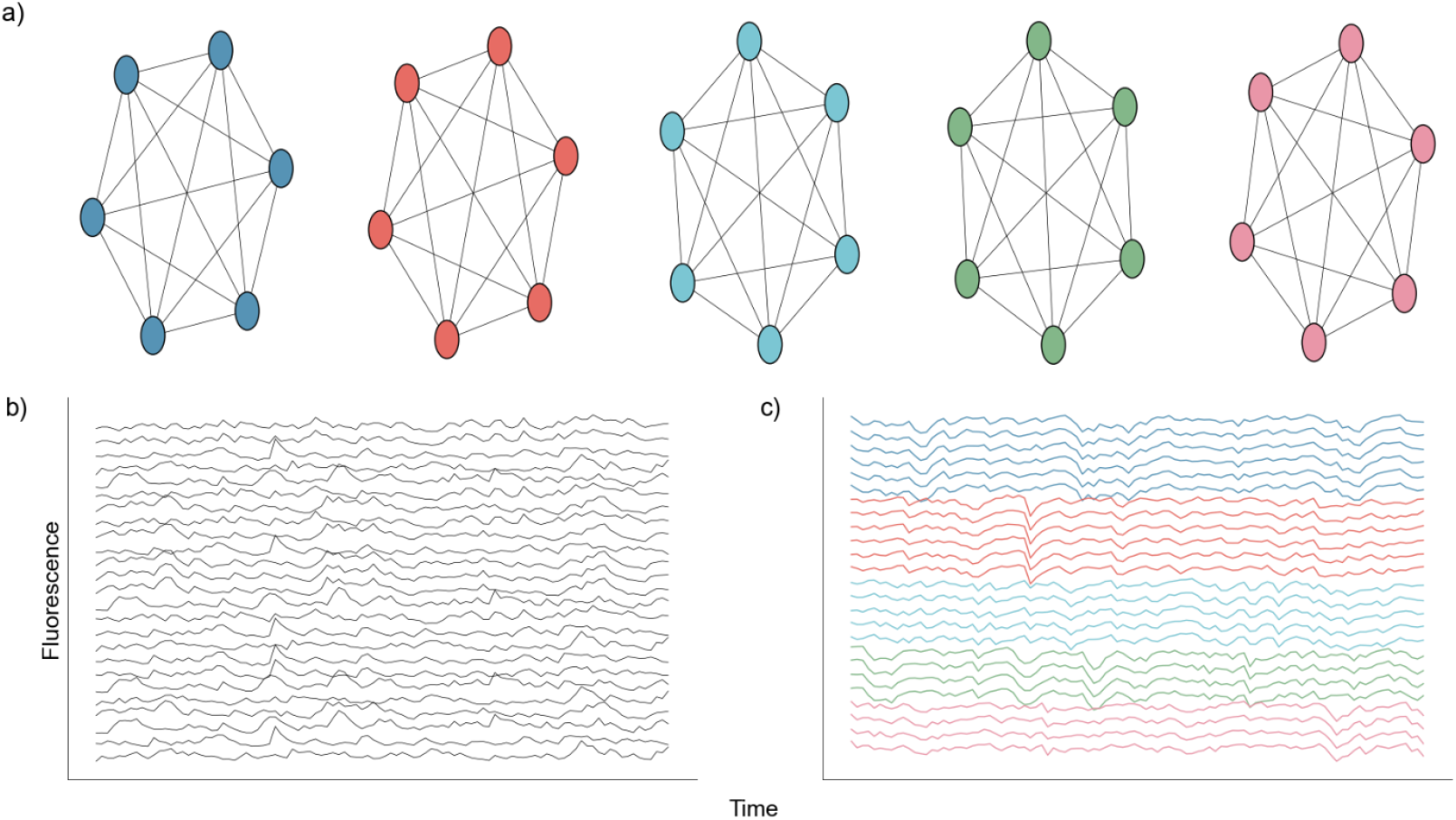
Nodal Network Architecture. **a)** Diagram showing perfect intraconnection between neurons and independence between nodes. **b)** Randomly selected calcium traces generated by the network. **c)** Same randomly selected calcium traces shown in b), but sorted according to node

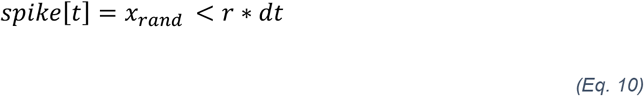

where a neuron spikes at time t if xrand (uniform [0, 1]) is less than the product of the set firing rate, r, and the time step, dt. We update this generator to include additional network effects, and a refractory period. We represent the connectivity between neurons as an adjacency matrix A. Within any node of 20 neurons, A is 1, while between nodes A is 0; we exclude self-connections by setting the diagonal of A to 0. We also include a refractory period of duration q time steps, such that after a spike a neuron is temporarily unable to generate a subsequent spike.

We simulate the spiking activity of a single neuron in this network n = {1,2, … N} as:

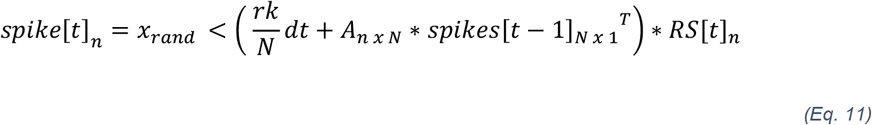

where r = 3 is the set nominal rate, N = 100 is the total number of neurons, k = 5 is the number of nodes, and dt is the time step. A is the adjacency matrix of dimensions N x N, where each row represents the incoming connections to a neuron, and each column represents the outgoing connections of a neuron, and *RS*[*t*]_*n*_ *=* {0,1} *i*s the refractory state of neuron n at time t. A neuron spikes at time t if the randomly generated number *x*_*rand*_ (uniform [0,1]) is less than the product in Eq. 11. We note that any multiplication of an adjacency of 1 to neuron n, in conjunction with a spike at time t-1 will result in a spike at time t in neuron n, given the neuron n is not in a refractory state. After simulating the network we convolve each spike train with a calcium kernel ^28,55^ as described below. The result is a series of 100 fluorescent neuronal activities (Fig. 2b), of which there are five groups of twenty that are perfectly correlated (Fig. 2c).

### Random Process Propagation

We generate a second simulation paradigm to more closely represent a physiologically relevant dynamic in calcium fluorescence data: a small number of independent random spiking patterns simultaneously driving calcium events in all neurons. To do so, we first simulate k = 5 independent spiking patterns as,

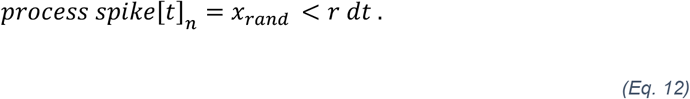

We then consider a population of N = 150 neurons that receive inputs from each of the underlying spiking patters (Fig. 3a) weighted as follow: 20% of the neurons have weights uniformly distributed between 0.2 and 1.0, and 80% of the neurons have weights exponentially distributed (rate parameter *λ = 8*.*0472) wi*th maximum value of 0.2 (Fig. 3b). In this way, most neurons are connected with weights following an exponential distribution ^56^, and the few neurons with strong weights drive a majority of the activity. The resulting adjacency matrix A between the k spiking patterns and N neurons has dimensions k x N, where each element describes the weight of the kth underlying process input to neuron n = {1, 2, … N}. We simulate each neuron’s spike train as:

**Fig. 3.**
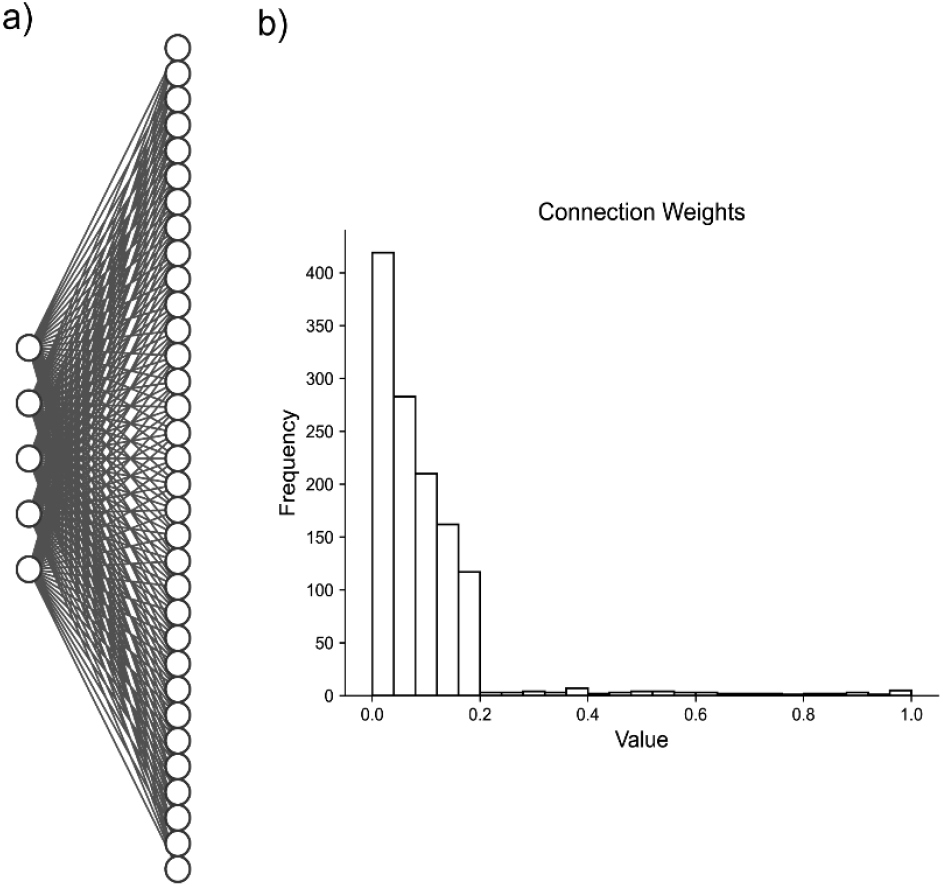
Artificial Random Propagating Process Network Architecture. **a)** Sample connection diagram showing all random processes projecting to all neurons. **b)** Single histogram of generated connections between processes and neurons from sampled distribution

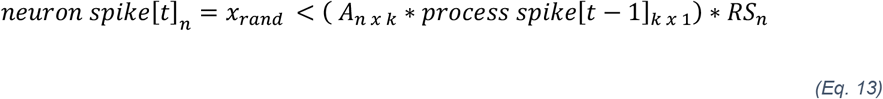

where n indicates the nth neuron, k indicates the kth underlying spike processes, and we include a refractory period for each neuron. Doing so results in N = 150 neuronal spike trains. We convolve the spike train of each neuron with the calcium kernel ^28,55^, resulting in simulated calcium data similar to our in vivo recordings, mirroring: activity rate, activity distribution, underlying macroscopic drivers, and calcium neuronal time series characteristics.

### Calcium Kernel

We convolve the simulated spike trains with the following calcium kernel to represent the fluorescence recorded from the genetically encoded calcium (GECI) kernel jGCaMP7f:

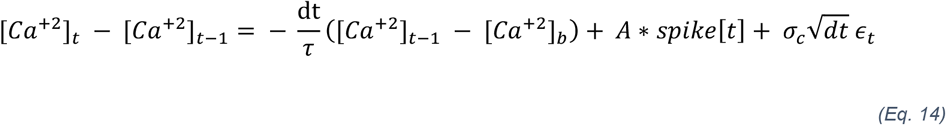

where 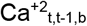 are the calcium concentrations at t, t-1, and the baseline, respectively. Here, dt is the time step, *τ i*s the time constant of the GECI, σc is the variance of the calcium noise, and ϵt is a random variable following the standard gaussian distribution. We then calculate the fluorescence as:

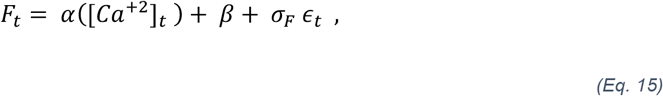

where Ft is the fluorescence at time t, α is the intensity, *β i*s the bias, σF is the variance of the calcium noise, and ϵt is a random variable following the standard gaussian distribution. We set *τ* to 0.265 and σc to 0.5 to match the kinetics of jGCaMP7f ^57^. We set the calcium baseline to 0.1, A and α to 5, β to 10, and σF to 1 ^28^.

### Component Assignment Success

Perfectly Connected, Intraconnected Nodes:

To assess whether nodes were successfully assigned to components, we develop a framework for each of our generated network architectures. For the nodal model, we begin by summing the weights for each node, in each of the decomposed components, described by the pseudocode below:

Input: H (matrix of neuronal weights), nodes (array detailing which neurons belong to each node) Output: aggregated_weight_matrix

1. aggregated_weight_matrix = zeros(num_nodes, num_components)
2. for i in range(num_nodes)
3. for j in range(num_compnents)
4. aggregated_weight_matrix [i, j] = sum(H [nodes[i], j)

The result is a five by five matrix, where the element [i, j] is the sum of the weights for neurons in the ith node for the jth component. To then determine success of prediction, we establish two conditions that must be met. First, within a particular component, the sum of weights for the node with the highest value should exceed the sums of weights for all other nodes within the same component. Second, this highest sum should also be the greatest for the node, across all components. By applying these conditions, we can assess the assignment for each node individually. In essence, once we obtain the aggregated weight matrix, an ideal model would exhibit the same element as the maximum value for each row and column.

Random Process Propagation:

We calculate a 5 x 5 correlation matrix between the weights in each of the 5 components and the connection probabilities from each of the 5 generated random processes sampled from the defined exponential distribution. To assess assignment, we again consider two conditions. First, for a chosen component, the highest correlation with the connection probabilities from one random process should surpass the correlation to the other random processes in the same component. Second, this highest correlation within each component should identify a different random process; i.e., each component should identify a unique random process. We consider a component successfully assigned to one of the five random processes if its highest correlation is more correlated with that process than the other four processes.

### Awake-State Data

To investigate the application of each DR technique to *in vivo* recordings, we also consider calcium imaging data recorded from 36 awake mice. Briefly, recordings in layer 2/3 of murine S1 ^58^ were collected during the awake, resting state ^8^. Traces representing calcium transients were extracted from the recordings and processed using standard techniques ^58^.

## Results

We test NMF against three existing DR approaches in common use: UMAP, PCA, and ICA. We do so using two different simulated neuronal network architectures to assess the relative performance of each DR approach, when the network structure is known. We first build a simple network with five groups of independent neurons, each with 20 neurons, in which the neurons in each independent node are perfectly connected (Fig. 2). In this architecture, a spike in one neuron in a node will drive all other neurons in that node to spike. We use such an architecture to test if, and how well, models can identify the five nodes given traces of simulated activity.

To further evaluate the performance between DR methods, we examine a more intricate network architecture aligned with our spontaneous calcium recordings. In doing so, we create five distinct random processes driving 150 neurons at varying degrees of strength (Fig. 3). This approach better reflects the dynamics we aim to capture, encompassing underlying activity patterns in diverse contexts driven by weighted sub-networks of neurons. Utilizing this surrogate architecture, our objective is to assess how effectively the models can extract both the underlying activity patterns, and the corresponding contributions by individual neurons, given a clear end target for performance assessment. We present two different analyses for both simulated network architectures. We both perform a qualitative analysis for a single generated network, and a more scaled quantitative approach for a series of 256 random instantiations and simulations of the networks. We find that NMF best captures the dynamics of the simulated networks, with components summing to provide an accurate macroscopic low dimensional representation of the data. We argue the parts-based representation provided inherently by NMF supports the improved performance for these data and the goal of interpretably extracting low dimensional neuronal network dynamics.

### Perfectly Intraconnected, Independent Nodes

We generate the nodal networks and fit each of the DR methods to the simulated calcium traces. Initially, we evaluate the average variance explained and the AIC across 256 networks to gauge how well the models capture the original data at different lower dimensionality levels. Both NMF and PCA exhibit similar variance explanation patterns across all components, with the average variance explained stabilizing at five components (Fig. 4a, left axis), as required to capture the dynamics of simulated activity in five nodes. This stabilization is influenced by the introduction of unique noise increments in each trace (Eq. 14 & Eq. 15). Additionally, the AIC for the NMF method consistently minimizes at five components across all 256 model instantiations, effectively optimizing for the correct number of nodes (Fig. 4a, right axis).

**Fig. 4.**
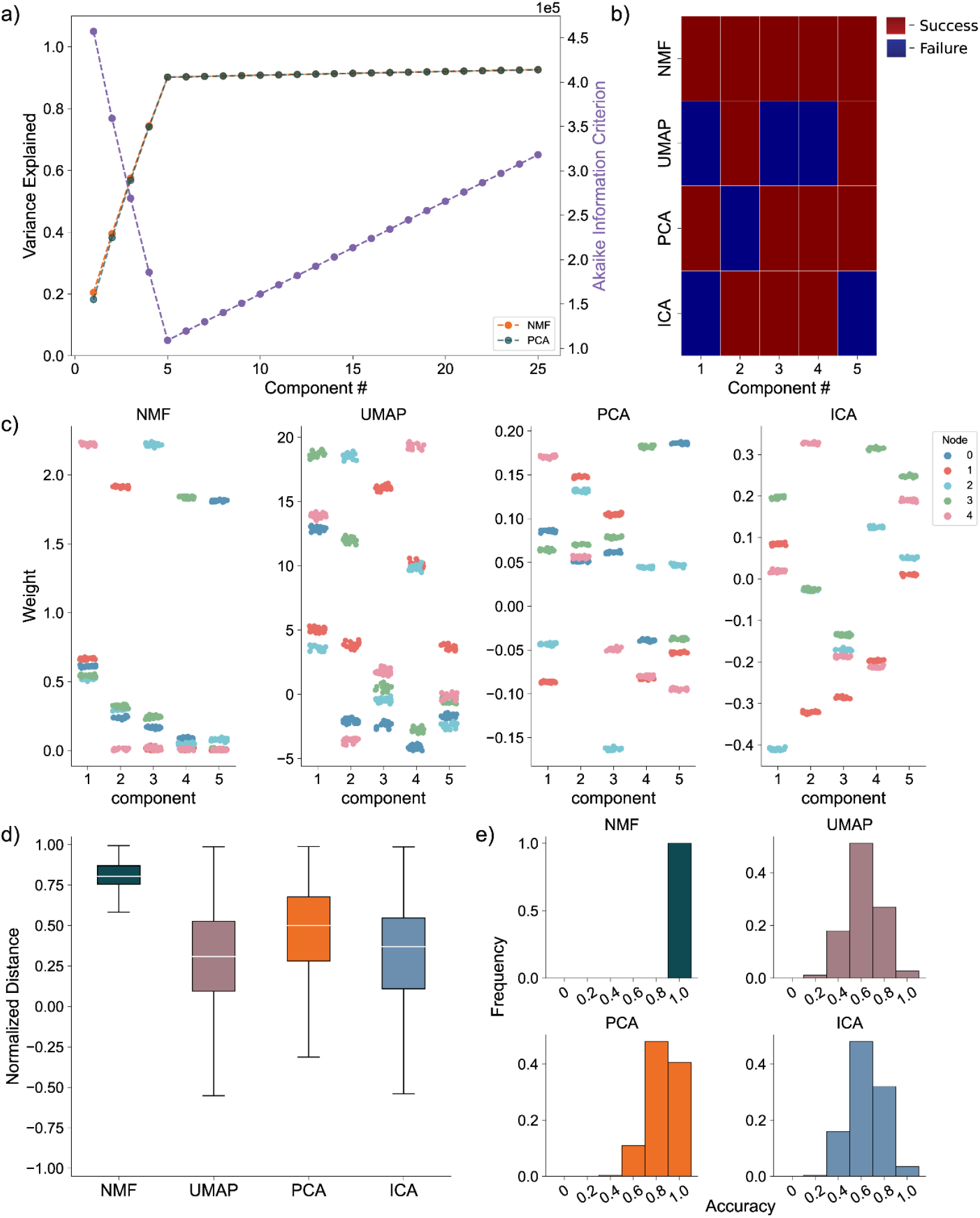
Artificial Nodal Network Analysis. **a)** Variance Explained for NMF and PCA, left axis (n = 256 networks) per component added to the model, and Akaike Information Criterion (AIC) for NMF, right axis (n = 256 networks). **b)** Prediction visualization between nodes and components for single network. **c)** Neuronal weights (y axis) for each model (panel) for each node (color) for each component (x axis). **d)** Average between neurons inside and outside predicted nodes for each component (n = 256 networks). **e)** Model assignment accuracy (n = 256 networks)

To evaluate how the different approaches capture the independent nodes of activity as components, we devised a discrete metric to determine success or failure of individual node assignment to components, detailed in the methods, for each architecture. We show the assignment success for a single model for each of the detailed DR methods (Fig. 4b). We find the best performance with the NMF method, which successfully maps each node to a component, followed by PCA, ICA, and UMAP. We repeat this process for 256 replicate models (Fig. 4e) and find that NMF always maps each node to a separate component, followed by PCA which performs with a mean accuracy of 0.8578, and then ICA and UMAP which perform with mean accuracies 0.6445 and 0.6242, respectively, and typically fail to successfully map the entire model.

We then visualize how each of the DR methods map the nodal patterns of activity to each of the components. We do so by showing the weight for each neuron, for each component, for each method, delimited by node (Fig. 4c), for a single instantiation of the simulated neural activity. In this example, we find a very clear separation of weights from each node for the NMF method; each component assigns weights to the neurons within a node. We argue this separation is a result of the parts-based representation that NMF inherently embodies ^35^. In a parts-based representation, each component contributes an additive combination of components to faithfully represent the original data. This means that NMF decomposes the overall activity into identifiable parts, enabling a clear separation of weights for each node. Conversely, the PCA and ICA methods produce distributions of node weights with less clear separations within a component. While the first 5 components of the PCA method each identify neurons in each node, the constraints of the method result in counterbalancing variables that obfuscate the underlying input resulting in the data, hindering performance ^37^. UMAP poorly separates nodes into five components because a strength of UMAP, and manifold learning in general (e.g., TSNE), is visualization in very low dimensions (restricted to two or three dimensions) ^24,25^.

To provide a quantitative assessment of model performance, we calculate the average displacement between neuronal weights assigned to a component and all other weights within the same component, for all components of each of the 256 instantiations of the model (Fig. 4d). We do so to provide a quantitative and continuous measurement of model performance (separation between weights in node and not in node, at assigned component) beyond binary success/failure, with the magnitude indicating the degree of separation between the neurons in the assigned node and all other neurons within a given component. Normalized per model, a positive value describes an assigned node whose average weight was higher than the weights of the other node in that component. A negative value would then describe the opposite, with the magnitude of the displacement describing the degree of difference between the weights. Our findings reveal NMF best separates nodes into components, followed by PCA, UMAP, and ICA. While this supports our finding that NMF specifically assigns patterns of nodal activity to components, the other methods do tend to separate a majority of the nodes of neurons individually in each component (the normalized distance tends to exceed 0), with the differing representations being attributable to the different mathematical constraining during fitting (UMAP – very low dimensional manifold, PCA – orthogonality, ICA – independence). However, as we show in our next case, this structure breaks down on the introduction of more complicated data.

### Random Process Propagation

We simulate a network with a more complicated architecture to further evaluate model performance. This architecture involves generating five underlying random processes to drive the network activity. Each underlying random process provides input to each of 150 neurons, with a weight randomly sampled from a generated exponential distribution (Fig. 3). By analyzing these artificial networks, we aim to probe how well NMF maps the random process by observations from the received neuronal inputs to components. We similarly aim to compare the performance and low dimensional representation of NMF and the other methods we implement.

Analyzing the average variance explained for all 256 uniquely seeded networks of this architecture, we find that NMF explains significantly more at all components (Fig. 5a, left axis). We argue this is a function of the difference between optimizations. Rather than having a strict orthogonality constraint between dimensions as in PCA, NMF aims to reconstruct the data with as little error as possible into the specified number of dimensions, allowing for explaining more variance with fewer components. The AIC shows a minimum at five components, the number of random processes used to generate the network (Fig. 5a, right axis).

**Fig. 5.**
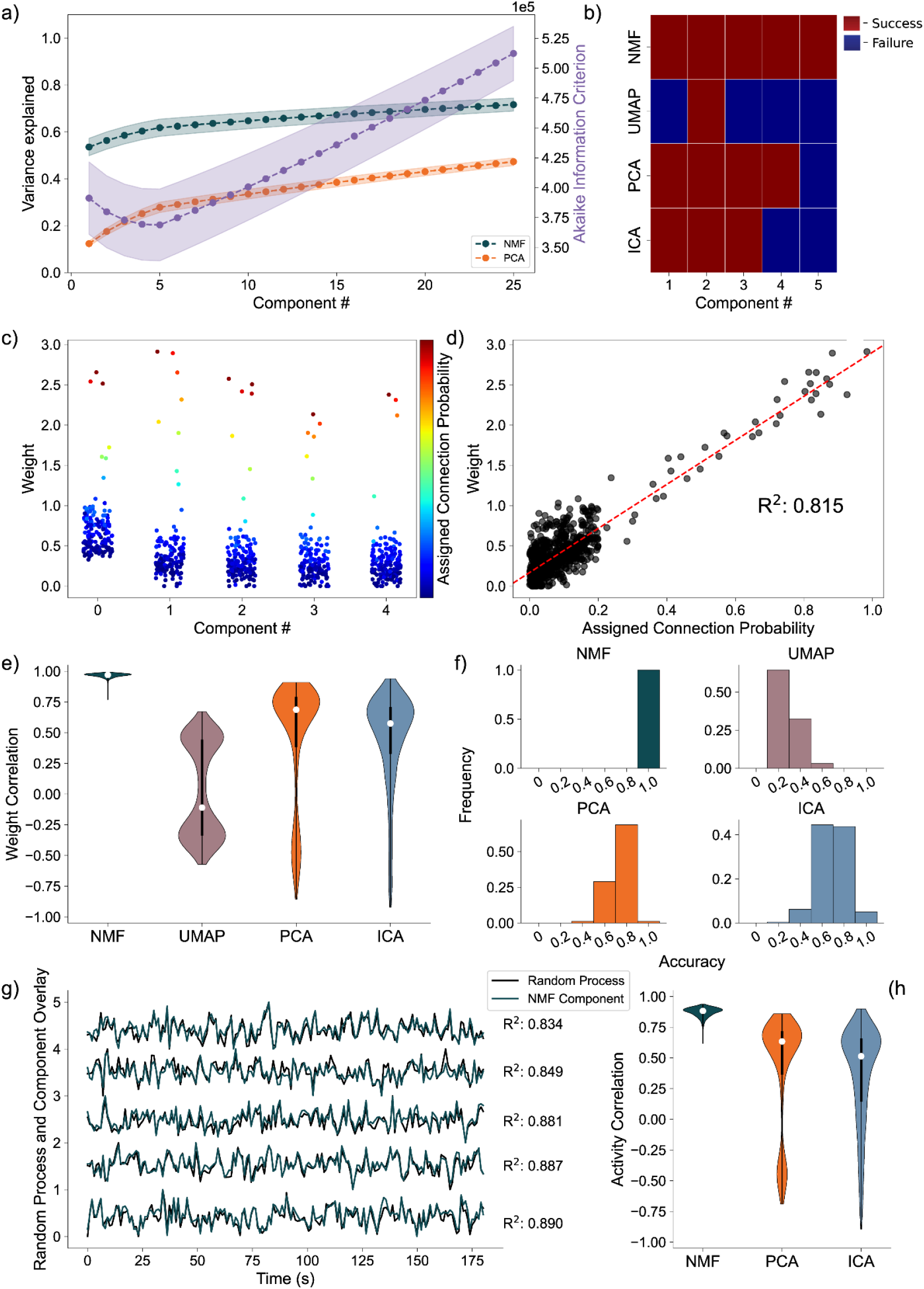
Artificial Random Propagating Process Network Analysis. **a)** Variance Explained for NMF and PCA, left axis (n = 256 networks) per component added to the model, and Akaike Information Criterion (AIC) for NMF, right axis (n = 256 networks). **b)** Assignment visualization between nodes and components for single network. **c)** Neuronal weights for NMF model (y axis) for each component (x axis) compared to assigned connection probabilities (color scale) for the predicted random process. **d)** Correlation between neuronal weights for NMF model (y axis) and the respective assigned connection probability (x axis). **e)** Average Correlation between neuronal weights for model and the respective assigned connection probabilities (n = 256 networks). **f)** Model assignment accuracy (n = 256 networks). **g)** Correlation between component activity from NMF model and assigned random process activity. **h)** Correlations between component activity from model and assigned random process activity (n = 256 networks)

We quantify assignment success of this model differently from the nodal architecture, due to the more continuous nature of the data, detailed in the methods. For or an example model instantiation (Fig. 5b), only the NMF method successfully assigns each process to one component. Assessing the performance over 256 realizations of the network (Fig 5f), we find that NMF successfully assigns each random process to a component. Consistent with the results for the nodal architecture, the other methods (PCA, ICA, and UMAP) have reduced performance. We conclude that NMF reconstructs the latent network structure from the observed data, capturing the five underlying random processes in a parts-based representation.

Beyond discrete prediction, we consider the correlation between the assigned connection probabilities from the latent processes to the observed neurons, and the neuronal weights of the predicted component. To start, we visualize the results for one instance of the NMF model, relating the neuronal weights of each component to the assigned outgoing connection probability from the assigned underlying random process to each neuron (Fig. 5c). We find that large neuronal weights in each component tend to occur for large connection probabilities for the predicted random process with a high correlation of 0.84 (Fig. 5d). Repeating this analysis for 256 realizations of the model, and regardless of prediction success, we find that NMF consistently infers components with high correlations between the connection probabilities and weights (Fig. 5e). The other methods have reduced performance, consistent with the results for the nodal architecture with PCA, ICA, and UMAP following in performance, respectively.

We finally analyze how the activity of each component correlates with the activity of the underlying random process. Qualitatively, for a single realization of the model, the component activity captures the underlying processes very well (Fig. 5g) with correlations between the activities exceeding 0.85. Repeating this analysis across 256 realizations of the network, we once again find that NMF best captures the activity of the latent input processes, followed by PCA and ICA (Fig. 5h); we note that UMAP does not estimate the times series of the latent processes.

We propose that the positivity and parts-based representation of the NMF method enable accurate reconstruct of the high-dimensional activity through low dimensional inputs. We propose that the constraints implemented by the PCA and ICA methods may obfuscate inputs to the observed neural network in its low dimensional representations. Finally, we interpret the poor performance of the UMAP method as indicating this method is better adapted for low dimensional visualization of less variant data. We conclude that NMF accurately reconstructs the latent inputs to a biophysically-motivated neuronal network that simulates calcium fluorescence recordings, despite multiple barriers to accurate identification (e.g., the underlying process is unobserved, with random connectivity to the observed neurons whose calcium signal is obfuscated with two types of noise). These results demonstrate NMF outperforms existing methods in common use to extract the underlying dynamics present in a series of neuronal activities recorded in calcium imaging.

### Awake-State Data

As a final proof of concept, we analyze application of the DR methods to awake state calcium imaging recordings from 36 mice. In doing so we apply the same analytical framework as for the simulated data. Analysis of the variance explained for each model (Fig. 6a) shows similar performance between the NMF and PCA methods, with a weak trend towards increased values for the NMF method. This small advantage in increased variance explained may be due to the NMF method acting to capture the original data as quickly as possible into a set number of components (Eq. 2), rather than using a complex geometrical constraint to analyze the data^50^. We further find that NMF, using AIC, identifies an optimal number of components to describe the data in each of the recordings (Fig. 6b), with number of neurons helping inform the number of components at a correlation of 0.662. We qualitatively characterize an NMF model fit to a single recording, using 18 components as determined by AIC. When characterizing the neuronal weights estimated by the NMF method, we find behavior consistent with the simulated random process network (Fig. 6c); the NMF method results in in the neuronal weights with an approximate exponential distribution (Fig. 6d), with most neurons contributing little to the overall network activity, and few neurons very significantly contributing. We further briefly characterize the correlations between the activity for each decomposed component (Fig. 6e), finding that pairwise correlations are consistently low between them (Fig. 6f). Given that the correlations are low, and we capture a significant portion of the variance, we interpret these results to indicate that the NMF, and the parts-based nature of the model, successfully extracts unique predominant patterns of activity in the data. We conclude that, applied to these *in vivo* calcium imaging recordings, that the NMF method identifies components with distinct (uncoupled) dynamics. While we demonstrate here that we successfully decompose statistically similar data from our simulations into statistically similar models, future work will relate inferred components and dynamics to experimentally elucidating biological mechanisms and behavior. We conclude that the NMF method is a promising tool to analyze neuronal network dynamics and identify meaningly sub-network activity via a relatively simple and interpretable approach.

**Fig. 6.**
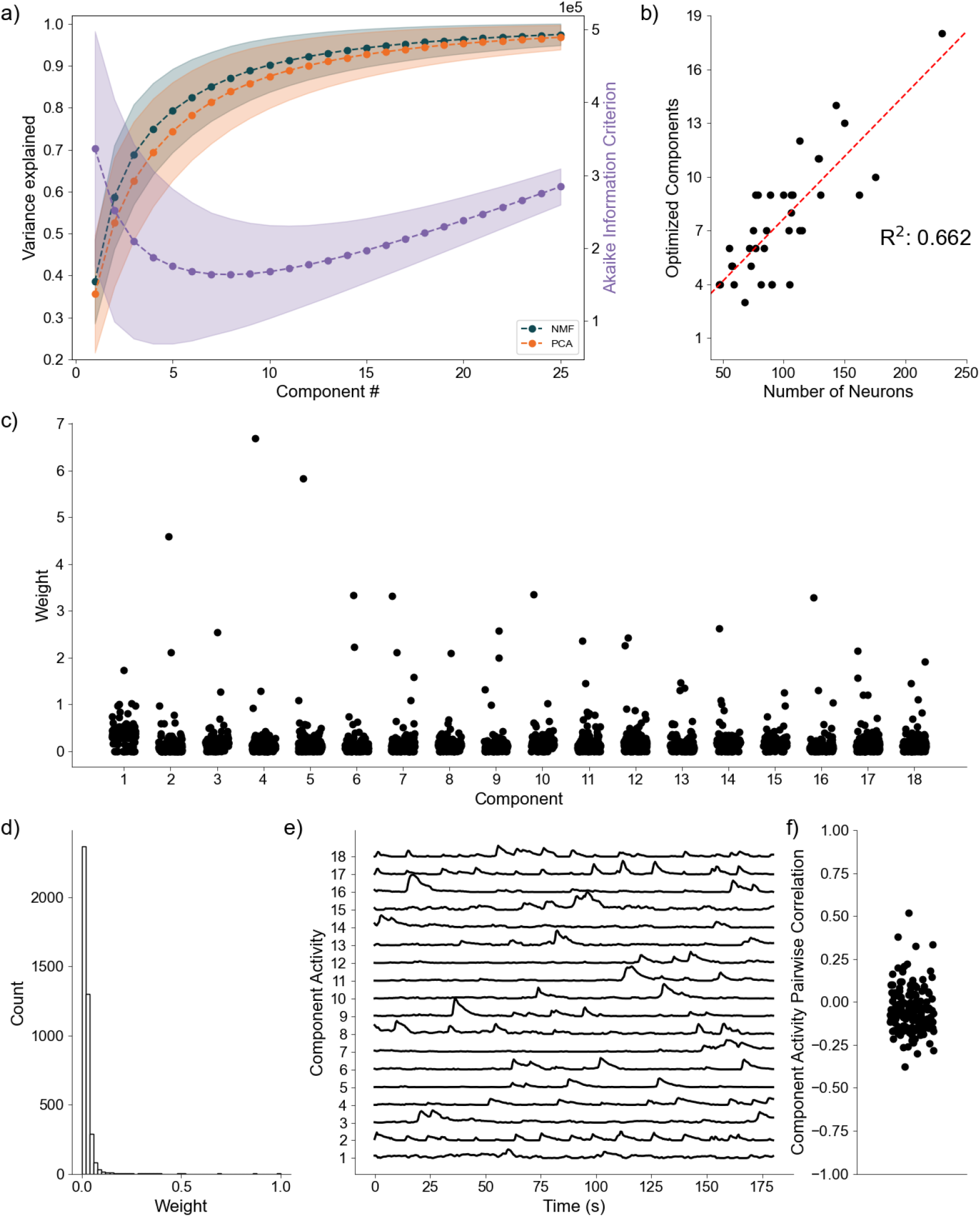
Baseline Awake State Network Analysis. **a)** Variance Explained for NMF and PCA, left axis (n = 36 mice) per component added to the model, and Akaike Information Criterion (AIC) for NMF, right axis (n = 36 mice). **b)** Correlation between number of neurons (x axis) and number of optimized components (y axis) (n = 36 mice). **c)** Neuronal weights for NMF model (y axis) for each component (x axis) for a single mouse. **d)** Neuronal weight histogram from NMF model for the same mouse as c). **e)** Component activity for NMF model of the same mouse as in c) and d). **f)** Pairwise correlations between component activities shown in e)

## Discussion

We have developed an analytical pipeline to more thoroughly model neuronal network dynamics with NMF, considering all information provided by the model in its low dimensional representation. We generate a series of calcium traces recorded from ground-truth artificial neuronal networks at two degrees of complexity to assess how well our analyses extract the ground-truth responsible for the simulated dynamics, and compare the performance of NMF to UMAP, PCA, and ICA. We find the NMF significantly performs the best, followed by PCA, ICA, and UMAP, respectively. We then apply our NMF pipeline to a series of in vivo baseline recordings and find that NMF confers a similar, and easily interpretable representation to the underlying random process network.

We argue this performance is a result of the “parts-based representation” conferred by NMF ^35^. Because every single element of the model is necessarily positive, and the decomposition is aiming to recapture the original data, every component sums to give a low dimensional macroscopic representation of the original data. Interpreting the decomposition as the sum of its parts, rather than a complex cancelling of variables to achieve an algebraic constraint, as in PCA ^37^, results in NMF being able to provide a better representation of the dynamics. Further, the only fundamental assumption made in NMF is that the data can be represented exclusively as positive values. Our other methods require more stringent assumptions. UMAP makes the assumption that the data can be embedded on a low-dimensional manifold^53^. While very advantageous in less variant data (large scale trial averaged electrophysiological recordings^52^), we argue our data is far too variant with limited scalability (single trial, single recording) for this method to be as effective. ICA assumes that components are both non-normal, and independent ^50^. While the assumption of non-normality of traces, variables, is a positive feature given that calcium traces are clearly non-normal data, we argue it is unfair to assume sub-network dynamics are completely independent of each other. Finally, PCA assumes the data is best described by the eigenvectors of the covariance matrix ^37^. We note that this algebraic constraint does provide PCA the unique attribute of the model being the same for the first i components, regardless of the number of components fit. NMF, UMAP, AND ICA will differ in their representation of the data, dependent on the number of components fit too. As a result, PCA is guaranteed to perfectly lossless in an inverse transformation of the data, given all components. However, PCA is clearly less efficient than NMF in generating compact descriptions of data like ours.

This pipeline was developed with the intent of analyzing state dependent neuronal network dynamics, and future work will analyze differential network dynamics that arise from unique experimental contexts. However, NMF, and other DR methods, require an underlying structure in the data and can be especially sensitive to potential extremes in the number of components and rates of activity in the network. An extreme sparsity or extreme saturation of events will lead to a much noisier representation in the model. Further, a fundamental assumption of DR is that the data can be represented with significantly fewer components than there are neurons ^9^. Data that would carry hundreds of underlying drivers of activity (or any case where the number of patterns approach the number of neurons) would be obfuscated in a model such as this. This analysis is able to go beyond summary statistics, providing a low dimensional representation of dynamics while still considering the activities of each neuron. Therefore, this approach will enable more sophisticated and nuanced analysis of different states of neuronal networks, and how they shift as a function of context.

In sum, NMF provides a superior method for the holistic analysis of network dynamics recorded in calcium imaging. The mathematical constraints required by NMF, linearity and positivity, complement the nature of fluorescent recording and neuronal activities well. Further, the parts-based nature of NMF provides a simple and interpretable representation of sub-networks of activity summing to drive macroscopic dynamics. As a result, we have developed an NMF pipeline to be an exceptionally valuable tool for elegantly demystifying shifting neuronal network dynamics.

## Acknowledgements

We thank Professor Chandramouli Chandrasekaran for providing a significant amount of the motivation for this research. We further thank Dr. Jacob F. Norman and Dr. Fernando R. Fernandez for their feedback and guidance in the development of this research.

## Data Availability Declaration

Code to simulate synthetic neuronal networks is available in python, and all statistics of all models fit to the simulated neuronal networks are available in python .pkl format (Fig. 4 & Fig. 5). The extracted activity of the 36 awake baseline recordings, and all optimum models fit to them are further available in python .pkl format (Fig. 6). All materials can be requested by contacting the corresponding author, John A. White, and will be shared in their entirety upon request.

## Author Contributions

Conceptualization, all authors; Methodology, DC; Validation, DC; Formal Analysis, DC; Investigation, DC; Resources, DC and JN; Software, DC; Data Curation, DC and JN; Writing – Original Draft, DC; Writing – Review and Editing, all authors; Visualization, DC; Supervision, JN, MAK, and JAW; Project Administration, DC; Funding Acquisition, DC and JAW.

